# Predicting brain atrophy from tau pathology: A summary of clinical findings and their translation into personalized models

**DOI:** 10.1101/2021.09.20.461165

**Authors:** Amelie Schäfer, Pavanjit Chaggar, Travis B. Thompson, Alain Goriely, Ellen Kuhl, for the Alzheimer’s Disease Neuroimaging Initiative

## Abstract

For more than 25 years, the amyloid hypothesis-the paradigm that amyloid is the primary cause of Alzheimer’s disease-has dominated the Alzheimer’s community. Now, increasing evidence suggests that tissue atrophy and cognitive decline in Alzheimer’s disease are more closely linked to the amount and location of misfolded tau protein than to amyloid plaques. However, the precise correlation between tau pathology and tissue atrophy remains unknown. Here we integrate multiphysics modeling and Bayesian inference to create personalized tau-atrophy models using longitudinal clinical images from the the Alzheimer’s Disease Neuroimaging Initiative. For each subject, we infer three personalized parameters, the tau misfolding rate, the tau transport coefficient, and the tau-induced atrophy rate from four consecutive annual tau positron emission tomography scans and structural magnetic resonance images. Strikingly, the tau-induced atrophy coefficient of 0.13/year (95% CI: 0.097-0.189) was fairly consistent across all subjects suggesting a strong correlation between tau pathology and tissue atrophy. Our personalized whole brain atrophy rates of 0.68-1.68%/year (95% CI: 0.5-2.0) are elevated compared to healthy subjects and agree well with the atrophy rates of 1-3%/year reported for Alzheimer’s patients in the literature. Once comprehensively calibrated with a larger set of longitudinal images, our model has the potential to serve as a diagnostic and predictive tool to estimate future atrophy progression from clinical tau images on a personalized basis.

## 1 Introduction

Alzheimer’s disease is associated with the production and propagation of misfolded oligomers of amyloid-*β* and tau proteins which have been found to accumulate in the form of amyloid-*β* plaques and tau neurofibrillary tangles in the brains of patients [1]. The early amyloid cascade hypothesis posits that amyloid aggregates are the primary catalyst of Alzheimer’s disease and its related cognitive decline [2, 3, 4]. In recent years however, the focus has shifted to hyperphosphorylated tau protein as a main agent in disease progression, as multiple studies have found a direct correlation between the amount and distribution of misfolded tau protein and cognitive impairment in Alzheimer’s patients [5, 6, 7, 8, 9]. Similarly, there appears to be a strong association of pathological tau with tissue atrophy. Tau pathology and cortical atrophy have been observed to follow the same stereotypical spatiotemporal pattern [10, 11, 12, 13], and tau positron emission tomography (PET) signal has been found to be a more reliable predictor for atrophy than either amyloid PET or baseline cortical thickness [8].

Mathematical modeling represents a promising, non-invasive means of investigating the mechanisms and characteristics of neurodegenerative diseases. Due to its direct correlation with tissue atrophy and cognitive symptoms in Alzheimer’s disease, tau pathology is a specifically interesting target for modeling, as it bears high potential for the prediction of disease progression timelines and the evaluation of potential future treatments. Several studies have focused on models for tau pathology and their validation with cross-sectional or longitudinal patient data [14, 15, 16]. We have previously proposed a network diffusion model to simulate misfolded tau protein propagation and shown that it can capture individual patient pathology by learning its model parameters from tau PET data of 76 subjects using Bayesian inference [17]. Similarly, several mathematical models have been used to shed light on the potential interplay between atrophy dynamics and the prionlike progression of misfolded tau deposition [18, 19, 20, 21]. Yet, to date, there is no physics-based model that integrates atrophy dynamics and Bayesian learning with multimodal neuroimaging data of both tau pathology and tissue atrophy.

Here we focus on two primary objectives: First, we conduct a brief review of atrophy dynamics in Alzheimer’s disease which motivates our physics-based network diffusion model of coupled tau progression and atrophy evolution. Second, we perform an inaugural Bayesian inference study to personalize its tau and atrophy parameters. This study lays the foundation for larger, predictive network atrophy studies and for the further development of more complex coupled tau and atrophy models. The remainder of the manuscript is organized as follows: In Section 2, we review literature on how atrophy manifests in Alzheimer’s disease and establish a link between the progression of atrophy and the spatiotemporal evolution of misfolded tau protein. Section 3 introduces a network proteopathy model augmented with an atrophy model motivated by the literature. In Section 4, we present experimental tau and atrophy data for four subjects from the Alzheimer’s Disease Neuroimaging Initiative [22] and demonstrate how to personalize the model using a probabilistic Bayesian approach. We discuss the features of our model in Sections 5 and 6, and offer concluding remarks and practical perspectives in Section 7.

## 2 Brain atrophy in Alzheimer’s disease

In this section, we summarize the characteristic clinical features of atrophy in Alzheimer’s disease, its temporal evolution, and its correlation to hyperphosphorylated tau protein. These observations lay the foundation for our physics-based models that couple the spatiotemporal evolution of tau pathology and tissue atrophy in Section 3.

### 2.1 Characteristics of atrophy

Tissue atrophy is a classical hallmark of neurodegenerative diseases. In Alzheimer’s disease, atrophy is regionally heterogeneous and presents in the form of volume loss, morphological changes, and cortical thinning in the gray matter of the brain [23, 11, 13, 24, 25], typically accompanied by white matter lesions [26], white matter tract changes [27, 28], and enlargement of the lateral ventricles [29]. The stereotypical spatiotemporal sequence of atrophy changes in the neocortex aligns with the topographic pattern of neurofibrillary tau tangles [11, 10]. Usually, volume loss is first observed in the medial temporal lobe, including the hippocampus and the entorhinal cortex, two structures whose volume changes are established biomarkers for Alzheimer’s disease progression [30]. With advancing disease, an increasing number of neocortical regions is affected by atrophy, initially the lateral temporal lobe, subsequently the parietal and frontal lobes. The sensorimotor and visual cortices are typically spared from atrophy until late in the disease [12]. Figure 1 summarizes and illustrates the most common characteristics of atrophy in Alzheimer’s disease: volume loss in the hippocampus, cortical thinning, morphological changes like the widening of cortical sulci, and the enlargement of the lateral ventricles as a consequence of a volume loss in adjacent gray matter structures.

**Figure 1:**
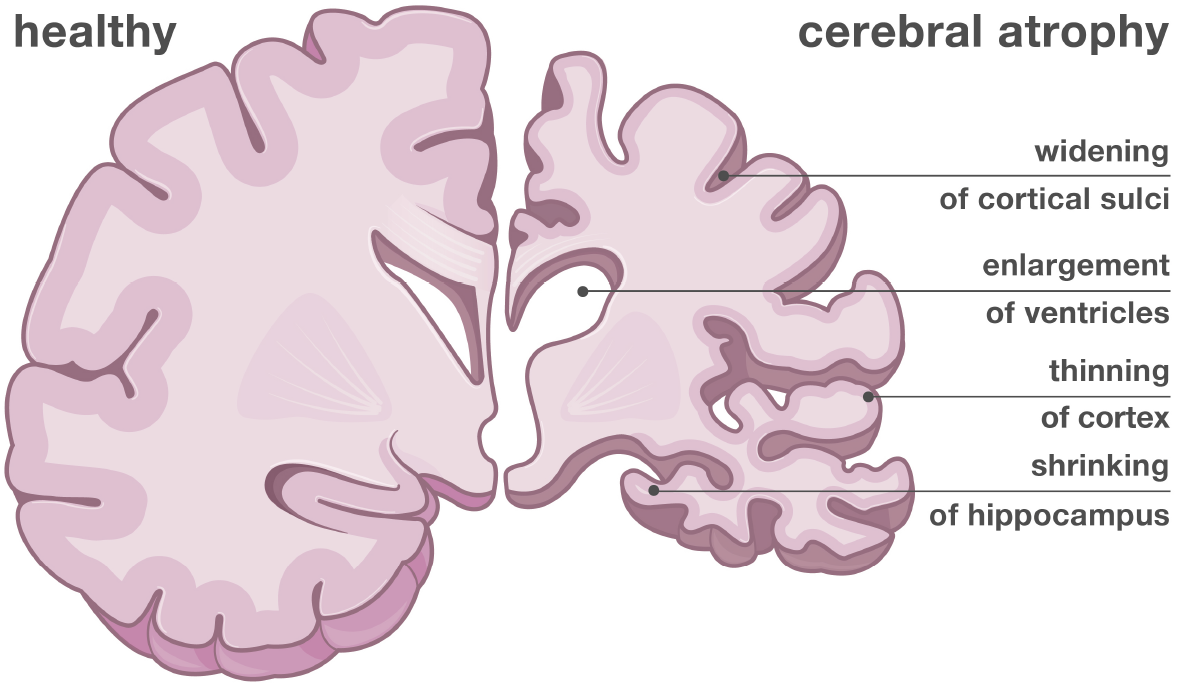
Atrophy in Alzheimer’s disease. Characteristic features of neurodegenerative atrophy in Alzheimer’s disease.

### 2.2 Dynamics of atrophy

Numerous brain imaging studies have investigated the characteristics of atrophy evolution in the pre-clinical and clinical phases of Alzheimer’s disease in comparison to healthy aging. These studies are commonly based on longitudinal magnetic resonance imaging (MRI) of a cohort of subjects combined with longitudinal cognitive assessments. Regional and global brain volume changes are evaluated from serial MRIs using a multitude of automated and semi-automated algorithms for image registration, segmentation, and morphometry.

#### Atrophy rates are higher in pre-symptomatic and symptomatic Alzheimer’s patients than in healthy controls

Throughout the course of a lifespan, all human brains exhibit a certain amount of tissue atrophy [31, 32, 33]. Increased cerebral atrophy is a widely-accepted biomarker for Alzheimer’s disease. Patients diagnosed with Alzheimer’s disease or mild cognitive impairment (MCI) show elevated rates of cortical thinning and volume loss in the hippocampus and across the whole brain compared to cognitively normal older adults [34, 35]. A study that compared hippocampal atrophy rates in mutation carriers for familial Alzheimer’s disease and non-carriers found that atrophy rates were significantly higher in mutation carriers, even before the onset of cognitive symptoms, and that the difference between the two groups increased over time [36]. Similarly, in a cohort of 240 subjects, whole brain atrophy rates were elevated for pre-symptomatic subjects who were later diagnosed with Alzheimer’s disease within six years after conclusion of the image acquisition [37].

#### Atrophy rates accelerate with disease progression

A commonly observed characteristic of Alzheimer related atrophy is an accelerated volume loss over the course of the disease. For example, whole brain atrophy rates and rates of ventricular enlargement were found to increase as patients progress from MCI to Alzheimer’s disease [38, 39]. Whole brain atrophy rates were found to accelerate with age even in cognitively normal controls, but acceleration increased by a factor of 1.5 in Alzheimer’s patients [40]. Two potentially intertwined explanations have been proposed for these accelerated whole brain atrophy rates: global acceleration could either stem from true acceleration of regional atrophy rates in already affected brain areas, or from the successive involvement of more brain regions as the disease progresses [38]. There is evidence of regional atrophy rates accelerating locally with disease severity when volume change is evaluated in different lobes [41] or even individual neocortical regions [42]. While atrophy rates appear to increase in all cortical regions with progressing cognitive impairment, the extent of local acceleration differs across areas [42].

#### Atrophy rates may decelerate at late disease stages

Observations of accelerating atrophy rates in Alzheimer’s disease are ubiquitous, but some evidence points towards decelerating atrophy rates beyond a characteristic inflection point. In a study that compared global cortical thinning rates in cognitively normal, MCI, and Alzheimer’s subjects, results indicate that atrophy rates first accelerate, but then decelerate with decreasing Mini Mental State Exam score. Even though data points from the low end of exam scores are sparse, the authors conclude that tissue atrophy follows a sigmoid like curve over the course of Alzheimer’s disease with an inflection point around an Mini Mental State Exam score of 21 [34]. Another study identified decelerating hippocampal atrophy rates around five years before death while atrophy rates in middle frontal and inferior temporal lobes continued to accelerate until death [43]. A similar study found that atrophy rates in the middle temporal lobe, while increasing significantly from cognitively normal to mild cognitive impairment, did not increase further with transition to Alzheimer’s disease. The study concludes that the middle temporal lobe might reach maximum atrophy rates already during mild cognitive impairment [41].

#### Alzheimer’s related atrophy rates are higher in patients with early disease onset

In paradox to the general trend that atrophy rates tend to increase with age, imaging studies have consistently identified higher atrophy rates for younger symptomatic patients than for older patients [44, 38, 45]. This is consistent with the theory that Alzheimer’s disease progresses more aggressively in patients with early disease onset [46, 47]. Table 1 summarizes the observed regional and whole brain atrophy rates from multiple clinical studies. Most of the included studies offer a comparison between Alzheimer’s patients at various stages of disease and cognitively normal, age-matched controls, providing evidence for increased atrophy rates in Alzheimer’s disease and regional acceleration of atrophy rates with disease progression.

**Table 1:** Atrophy rates. Reported whole brain and regional atrophy rates. Key: n, cohort size; C, controls; AD, Alzheimer’s disease; AD-EO, early-onset/familial AD; MCI-S, stable MCI; MCI-P, MCI progressing to AD; C-S, stable controls; C-MCI, controls progressing to MCI; AD-fast, fast progressing AD; AD-slow, slow progressing AD.

| study | subjects |  | atrophy rate in mean (SD)<br>%/year unless specified | other observations |
| --- | --- | --- | --- | --- |
| whole brain |  |  |  |  |
| Chan et al. (2003) [48] | AD-EO | n=12 | 2.8 (95% CI 2.3-3.3) | annual volume loss rises by 0.32 %/year<sup>2</sup> |
| Ridha et al. (2006) [36] | C-S | n=25 | 0.0 (0.57) | atrophy rates significantly higher in mutation carriers compared to controls, discrepancy increases over time |
|  | C-MCI | n=5 | 0.52 (0.51) |  |
|  | MCI-P | n=6 | 1.82 (0.87) |  |
|  | AD-EO | n=5 | 1.79 (0.74) |  |
| Sluimer et al. (2008) [45] | AD-EO | n=26 | 2.2 (1.1) | higher atrophy rates in early-onset AD |
|  | AD | n=39 | 1.7 (0.8) |  |
| Jack et al. (2004) [49] | C | n=40 | 0.4 (0.3) | atrophy rates are higher in subjects who progress from C to MCI than in non-progressors, and higher in AD with fast cognitive decline than in subjects with slow cognitive decline |
|  | C-MCI | n=15 | 0.8 (0.5) |  |
|  | AD-slow | n=32 | 0.6 (0.7) |  |
|  | AD-fast | n=33 | 1.4 (1.1) |  |
| Henneman et al. (2009) [39] | C | n=34 | 0.6 (0.6) | significant difference in atrophy rates between groups (C, MCI, AD) |
|  | MCI | n=44 | 1.3 (0.9) |  |
|  | AD | n=64 | 1.9 (0.9) |  |
| Leung et al. (2013) [44] | C | n=205 | 0.63 (95% CI 0.53-0.72) | acceleration in atrophy rates between groups (C, MCI, AD) but not evidence of acceleration within groups |
|  | MCI | n=352 | 1.01 (95% CI 0.92-1.1) |  |
|  | AD | n=156 | 1.41 (95% CI 1.23-1.59) |  |
| hippocampus |  |  |  |  |
| Barnes et al. (2008) [50] | C | n=212 | 1.41 (95% CI 0.52-2.3) | meta-analysis of nine studies on hippocampal atrophy rates |
|  | AD | n=595 | 4.66 (95% CI 3.92-5.4) |  |
| Henneman et al. (2009) [39] | C | n=34 | 2.2 (1.4) | significant difference in atrophy rates between groups (C, MCI, AD) |
|  | MCI | n=44 | 3.8 (1.2) |  |
|  | AD | n=64 | 4.0 (1.2) |  |
| Leung et al. (2013) [44] | C | n=205 | 1.22 (95% CI 0.9-1.55) | hippocampal atrophy rates accelerate by 0.22 %/year<sup>2</sup> |
|  | MCI | n=352 | 3.20 (95% CI 2.82-3.57) |  |
|  | AD | n=156 | 5.19 (95% CI 4.44-5.94) |  |
| Jack et al. (2004) [49] | C | n=40 | 1.4 (1.2) | atrophy rates are higher in subjects who progress from C to MCI than in non-progressors, and higher in AD with fast cognitive decline than in subjects with slow cognitive decline |
|  | C-MCI | n=15 | 3.3 (2.4) |  |
|  | AD-slow | n=32 | 3.0 (4.5) |  |
|  | AD-fast | n=33 | 3.6 (3.2) |  |
| McDonald et al. (2009) [42] | C | n=137 | 0.86 (1.27) | atrophy rates in all neocortical regions accelerate with clinical impairment in AD, atrophy is not linear or uniform across regions |
|  | MCI | n=231 | 2.39 (2.26) |  |
|  | AD | n=104 | 3.64 (2.12) |  |
| entorhinal cortex |  |  |  |  |
| Jack et al. (2004) [49] | C | n=40 | 2.9 (2.6) |  |
|  | C-MCI | n=15 | 5.1 (5.1) |  |
|  | AD-slow | n=32 | 8.0 (5.8) |  |
|  | AD-fast | n=33 | 8.4 (9.2) |  |
| Miller et al. (2013) [51] | C | n=81 | 1.0 (3.3) |  |
|  | C-MCI | n=20 | 2.7 (3.1) |  |
| McDonald et al. (2009) [42] | C | n=137 | 0.52 (1.13) | atrophy rates in all neocortical regions accelerate with clinical impairment in AD, atrophy is not linear or uniform across regions |
|  | MCI | n=231 | 1.93 (1.56) |  |
|  | AD | n=104 | 2.50 (1.13) |  |
| medial temporal lobe |  |  |  |  |
| Sluimer et al. (2009) [41] | C | n=34 | 0.6 (0.7) | atrophy rate in medial temporal lobe already reaches maximum before transition from MCI to AD |
|  | MCI | n=44 | 1.5 (0.7) |  |
|  | AD | n=64 | 1.5 (0.7) |  |
| Rusinek et al. (2004) [52] | C | n=21 | 0.37 (0.34) | annual atrophy rate increases with disease progression in AD |
|  | AD | n=18 | left: 2.2, accel. 0.73 %/year<sup>2</sup><br>right: 1.8, accel. 0.34 %/year<sup>2</sup> |  |
| ventricular expansion |  |  |  |  |
| Jack et al. (2004) [49] | C | n=40 | 1.7 (0.9) | expansion rates are higher in subjects who progress from C to MCI than in non-progressors, and higher in AD with fast cognitive decline than in subjects with slow cognitive decline |
|  | C-MCI | n=15 | 3.4 (1.6) |  |
|  | AD-slow | n=32 | 4.3 (3.3) |  |
|  | AD-fast | n=33 | 6.4 (3.7) |  |
| Leung et al. (2013) [44] | C | n=205 | 1.34 mL/year (95% CI 1.09-1.6) | rate of ventricular expansion accelerates in MCI and AD subjects |
|  | MCI | n=352 | 2.69 mL/year (95% CI 2.38-3.01) |  |
|  | AD | n=156 | 4.04 mL/year (95% CI 3.34-4.75) |  |

### 2.3 Evidence relating atrophy to tau pathology

Increasing evidence suggests that there is a close correlation between tau pathology and atrophy dynamics. In a study that combined longitudinal structural MRI within four years of death with postmortem neurofibrillary tangle autopsy, subjects with high tau burden showed higher gray matter loss in the medial and lateral temporal lobes compared to those with low tau burden [53]. Cross-sectional imaging studies have also shown that the pattern of neurodegeneration inferred from structural MRI mirrors the canonical progression of tau deposition in the brain [10, 6, 11]. Longitudinal imaging studies have consistently identified close correlations between regional tau PET signal intensity and a subsequent decrease in regional volume or cortical thickness in amyloid positive subjects and MCI or Alzheimer patients [54, 9, 55, 7]. A recent study based on baseline amyloid and tau PET scans as well as baseline and follow-up structural MRIs for 32 patients during the early clinical stages of Alzheimer’s disease, revealed the global tau PET intensity as a good predictor for future cortical thinning. Additionally, the results indicate a strong voxel-wise correlation between tau PET signal and future atrophy. Both on the global and local levels, tau PET was a stronger predictor for atrophy than amyloid PET or baseline cortical thickness [8].

## 3 A network model for tau pathology and atrophy

In this manuscript, we adapt the Fisher-Kolmogorov model [56] to characterize the local production and global spreading of misfolded tau protein across the brain. The continuous model is defined by

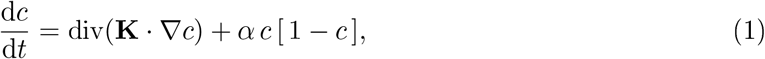

where *c* denotes the concentration of misfolded tau protein, and **K** denotes the diffusion tensor that contains information about the speed and directionality of protein propagation. The growth coefficient *α* captures how much misfolded protein is produced or cleared locally. We discretize the Fisher-Kolmogorov model on a network that represents the structural brain connectome. Suppose that *G* = { *E, N* } is the undirected graph of a structural brain connectome with edges *E* and nodes *N* that represent the anatomical regions of interest. A weighted adjacency matrix **W** is defined by the rule *W_ij_* = *W_ji_* > 0 whenever *n_i_* is connected to *n_j_* by an edge *e_ij_* in *E* and *W_ij_* = 0 otherwise. In this manuscript, we use an adjacency matrix constructed from diffusion tensor images of 426 participants of the human connectome project [57]. The network contains *N* = 83 nodes, each representing a cortical or subcortical brain region. Each edge *e_ij_* of the graph represents a tractographic approximation of the white matter connections between two regions of interest and is associated with a measure of the average fiber number *n_ij_* and fiber length *ℓ_ij_* along this connection. In line with a previous study [58], we define the weights of the adjacency matrix as 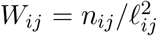. The files used to create the adjacency matrix are freely available [59, 60] as is the final adjacency matrix [61].

The weighted graph Laplacian **L** associated with the adjacency matrix **W** is then defined as

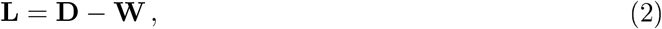

where **D** is a diagonal matrix defined by the row-sum of the weighted adjacency matrix as

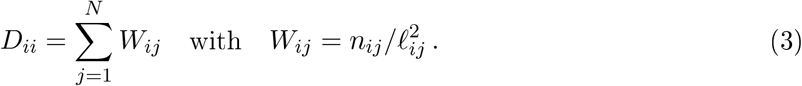

The Laplacian allows us to discretize the model on the network,

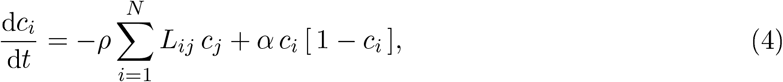

where the quantity *c_i_* denotes the normalized concentration of toxic tau protein in regions *i* = 1, 2,…,*N*, *ρ* is a transport coefficient, *α* is a growth coefficient, and *L_ij_* are the entries of the weighted graph Laplacian.

Motivated by the observations of Section 2, we couple the tau evolution model of Equation (4) to an atrophy model of the following form,

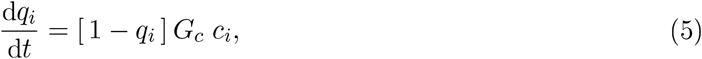

and express atrophy in terms of a regional relative atrophy marker 0 ≤ *q_i_* ≤ 1 for each node *i* = 1, 2,…, *N* in the connectome graph. The model postulates that the atrophy rate in a region of interest progresses based on the current atrophy level, *q_i_*, and the current amount of misfolded tau, *c_i_*, mitigated by a global tau-induced atrophy coefficient *G_c_*. The dynamics of Equation (5) capture several known features of the atrophy dynamics discussed in Section 2. The direct dependency of the atrophy rate on the nodal protein concentration *c_i_*, which itself follows a sigmoid curve over the course of the disease, allows us to capture the acceleration of atrophy rates observed in numerous longitudinal imaging studies. The saturation term [1 – *q_i_*] in Equation (5) ensures an asymptotic deceleration of atrophy rates towards the later stages of the disease.

## 4 Model personalization using medical images

Our coupled tau-atrophy model contains three global parameters: a global transport coefficient *ρ*, a growth coefficient *α* and a tau-induced atrophy coefficient *G_c_*. We have previously personalized the transport and growth coefficients *ρ* and *α* of Equation (4) using Bayesian inference and longitudinal tau PET data [17]. Here we also include structural information from magnetic resonance images to personalize the atrophy coefficients *G_c_*.

### 4.1 Subject data

We use longitudinal tau PET and structural imaging data from four subjects of the Alzheimer’s Disease Neuroimaging Initiative database [22] to prototype our model personalization using Bayesian inference. Two subjects were classified as cognitively normal (CN), two were classified as late mild cognitively impaired (LMCI). All four subjects were previously identified as amyloid positive [62]. All four subjects received four consecutive annual AV-1451 tau PET and T1 structural MRI scans, with an average time of 1.08 years between scans.

#### Tau data preparation

The processing of the tau AV1451-PET data by ADNI followed standard protocols [22, 63]. Briefly, for each subject, the PET images were co-registered to a corresponding high-resolution T1 weighted MRI, which had been segmented into 68 cortical and 45 subcortical regions according to the Desikan-Killiany atlas [64]. Each segmented region of interest corresponds to one node of our brain network from Section 3. For each region of interest, a PET standardized uptake value ratio was calculated using the inferior cerebellum as reference region. Since tau PET recordings in subcortical regions are known to be contaminated by off-target binding in the choroid plexus and nearby vascular structures [65, 66, 67], we exclude these regions from our analysis. To compare the clinically recorded PET tau values to our computationally simulated tau values, we map them into the range from zero to one. Briefly, we previously computed a regional tau positivity threshold of *c*^raw^ = 1.1 based on fitting a two-component Gaussian mixture model to the raw tau PET data from a larger cohort of 76 subjects [17]. We set values below this threshold to zero and normalize the remaining values with respect to the maximum occurring standardized uptake value ratio in our previous study, such that the normalized values *c*^pet^ range from zero to one, 0 ≤ *c*^pet^ ≤ 1. For each subject, we set the initial conditions for the protein field in our model to the tau uptake values measured in the baseline PET scan *p*^sim^(*t* = 0) = *p*^pet^(*t*_0_).

#### Atrophy data preparation

We process the structural data using FreeSurfer [68] via the Clinica [69] t1-freesurfer-longitudinal pipeline. This image processing pipeline consists of a sequence of different tools of the FreeSurfer software and involves the computation of a subject-dependent template space, segmentation of subcortical structures, extraction of cortical surfaces, parcellation of cortical regions, cortical thickness estimation, and the extraction of volume and thickness estimates in the template space at different points in time. We compute the volumes for all 83 cortical and subcortical brain regions that coincide with the 83 nodes of our brain network model from Section 3. We use the regional volumes measured at the baseline scan, 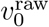, as reference points to which we normalize the regional volumes of all follow-up visits within each subject, 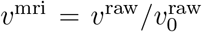. We then define the measured nodal atrophy as the relative reduction in volume, *q*^mri^ = 1 – *v*^mri^, with an initial atrophy value of 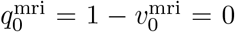. For each subject, we set the initial conditions for the atrophy field in our model to zero in all regions *q*^sim^(*t* = 0) = 0.

### 4.2 Bayesian inference

For each subject, we infer a personalized parameter set that most accurately explains the image data given the model in Equations (4) and (5). The parameter set 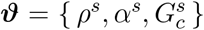 consists of a personalized transport coefficient *ρ*^s^, growth coefficient *α*^s^, and tau-induced atrophy coefficient 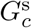. We approximate the posterior distribution 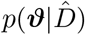 of the parameters ***ϑ*** given the imaging data 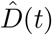 using Bayesian inference,

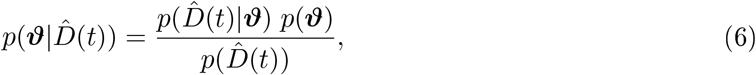

where 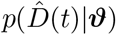 is the likelihood, *p*(***ϑ***) are the priors, and 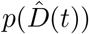 is the evidence.

For each subject, we specify a likelihood function that relates the computationally predicted tau and atrophy values at each region and time point, { *c*^sim^(*t_i_*), *q*^sim^(*t_i_*) } to the clinically recorded tau and atrophy values, { *c*^pet^(*t_i_*), *q*^mri^(*t_i_*) }. The time points *t_i_* for *i* = 0,1, 2, 3 are associated with the four consecutive years at which the PET and structural scans were recorded for each subject. To account for clinical measurement noise, we assume that the likelihood between the clinically recorded data 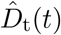 and 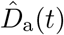 and the computationally predicted data *D*_t_(*t, **ϑ***) and *D*_a_(*t, **ϑ***),

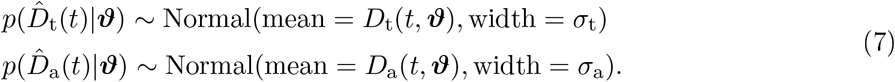

is normally distributed around the computationally predicted data with likelihood widths of *σ*_t_ for tau and *σ*_a_ for atrophy.

For the priors, we select weakly informative prior distributions for all model parameters in the set 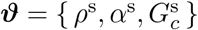 in addition to priors for the noise parameters that dictate the likelihood widths 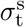 and 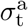. Table 2 summarizes the selection of our prior distributions.

**Table 2:** Prior distributions. Prior distributions for the personalized transport coefficient, growth coefficient, tau-induced atrophy coefficient, and the likelihood width.

| Parameter | Distribution |
| --- | --- |
| $\rho^s$ | TruncatedNormal($0 < \rho^s < 10$, mean=0, std=5) |
| $\alpha^s$ | Normal(mean=0, std=5) |
| $G_c^s$ | TruncatedNormal($0 < G_c^s < 10$, mean=0, std=5) |
| $\sigma_t^s, \sigma_a^s$ | InverseGamma(shape=2, scale=3) |

With the likelihood from Equation (7) and the prior distributions from Table 2, we compute the posterior distributions 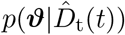 and 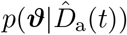 for the model parameters **ϑ** using Bayes’ theorem,

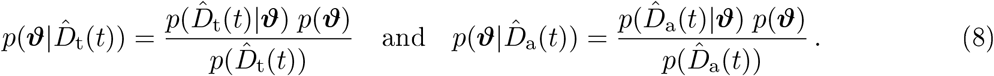

We adopt approximate inference techniques to numerically evaluate Equations (8) and personalize our model using the imaging data. We implement our model using two widely used Julia packages, the DifferentialEquations library and the Turing probabilistic programming library. The DifferentialEquations library [70] allows us to solve the differential equations (4) and (5) in time, to produce the model predictions *D*_t_(*t, **ϑ***) and *D*_a_(*t, **ϑ***). The Turing probabilistic programming library [71] serves to implement a Bayesian Markov Chain Monte Carlo sampling approach. It supports several different samplers from which we select the No-U-Turn sampler [72] to design a Hamiltonian Markov Chain Monte Carlo method. For each subject, we sample five chains with 1000 tuning samples and 5000 posterior samples per chain.

For each subject, we calculate the posterior model predictions and credible intervals for the global tau and atrophy evolution by propagating the uncertainty of each model parameter’s posterior distribution through our model. These predictions indicate how tau pathology and tissue atrophy may evolve over the next 30 years in each subject, based on the assumptions of our model and the information from the longitudinal imaging data.

## 5 Results

### 5.1 Imaging data

Figure 2 provides an overview of the regional volume data for our set of four subjects. For subjects 1, 3, and 4 we observe on average monotonically decreasing volumes over time. However, a number of regions seem to either exhibit an increase or no change in volume compared to the baseline scan. In all four subjects, we observe a positive correlation between regional tau load and the average atrophy in a region which agrees well with the assumptions based on which we constructed our coupled tau-atrophy model. This correlation is stronger in subjects 1,2, and 3 with Pearson correlation coefficients of R = 0.382, R = 0.397, and R = 0.494 respectively, than in subject 4 with R = 0.144. The regional volumes in subject 2 evolve less monotonically than those in the other subjects, and its average volume across all brain regions remains relatively constant over time. However, the plot shows that certain regions do decrease in volume compared to the baseline scan. Figure 3 and Figure 4 illustrate the collected tau and atrophy data on a template brain surface to show the regional distribution of the two fields in each subject.

**Figure 2:**
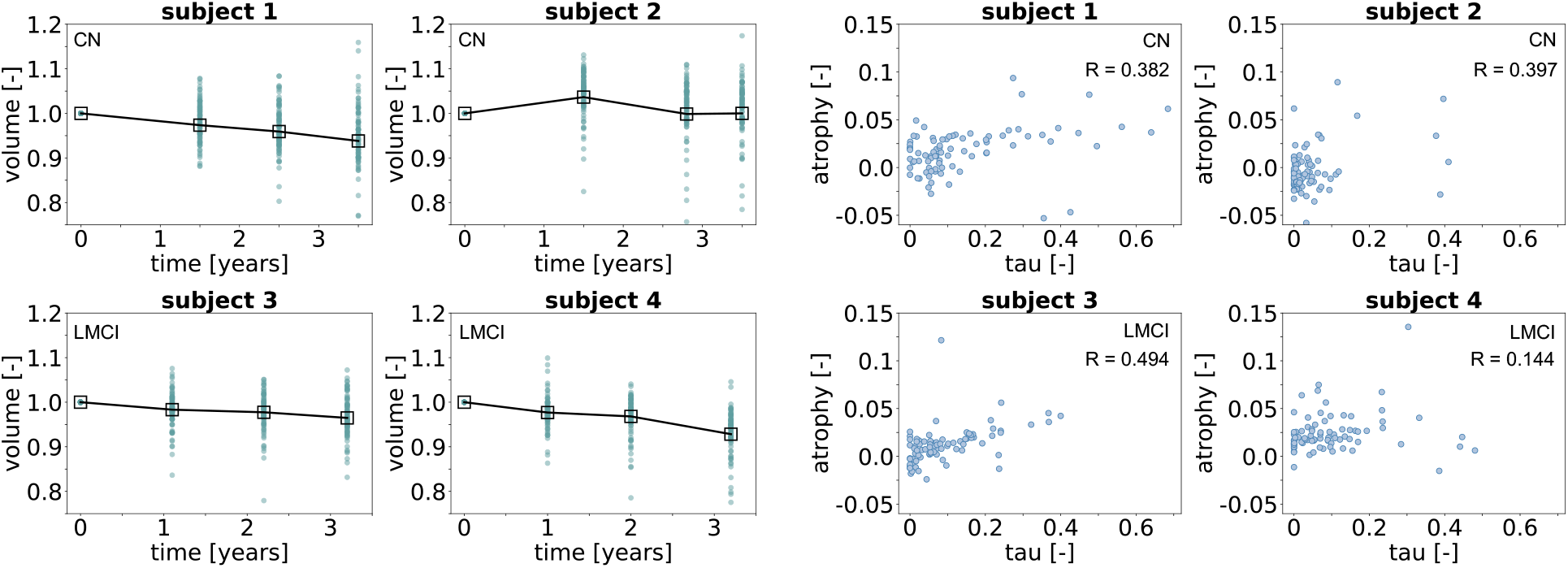
Subject data. Left: regional volume normalized to baseline over time. Each point represents one brain region at one visit. Squares represent the average volume across regions. Each subject’s clinical diagnosis is indicated in the upper left corner. Right: regional atrophy between visits averaged across all visits over regional tau load averaged across all visits. Each point represents one brain region. R indicates the Pearson correlation coefficient.

**Figure 3:**
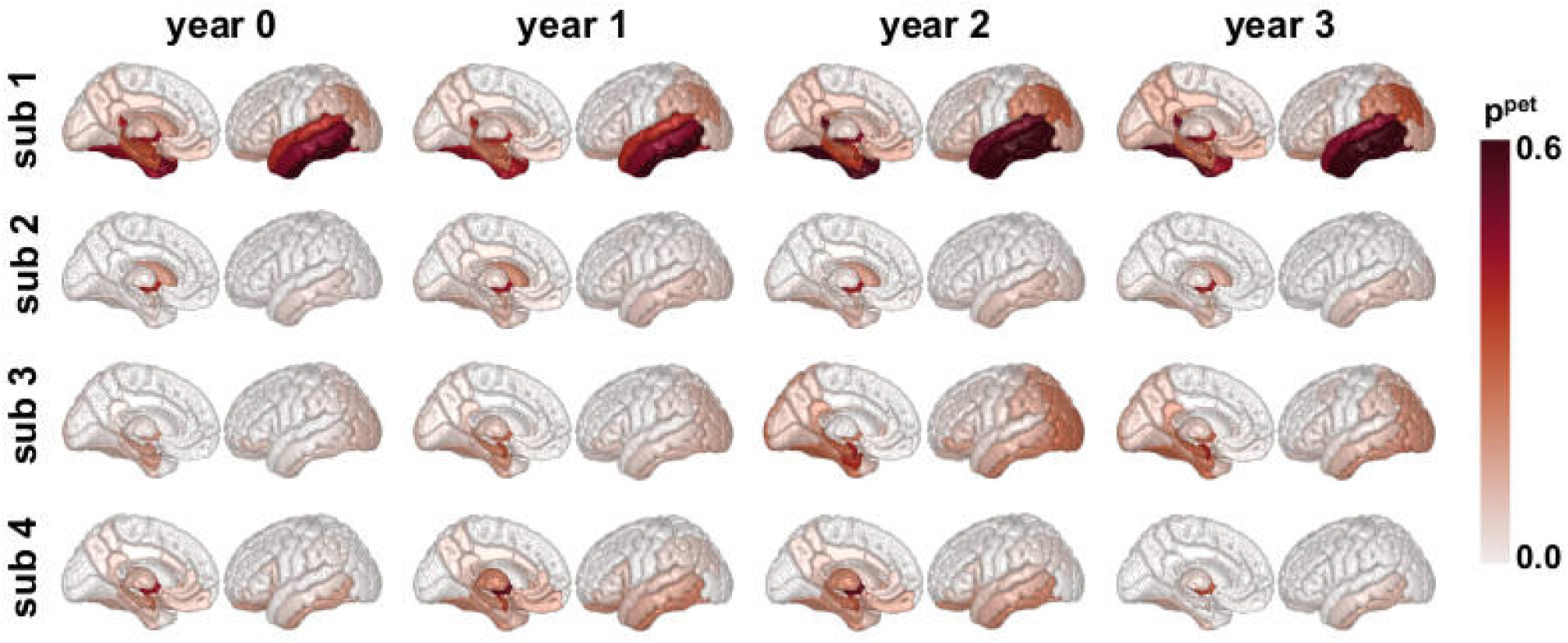
Subject data. Normalized misfolded tau concentration *c*^pet^ illustrated on a template brain. Each row summarizes the data for one subject, at baseline year 0 and three follow up scans.

**Figure 4:**
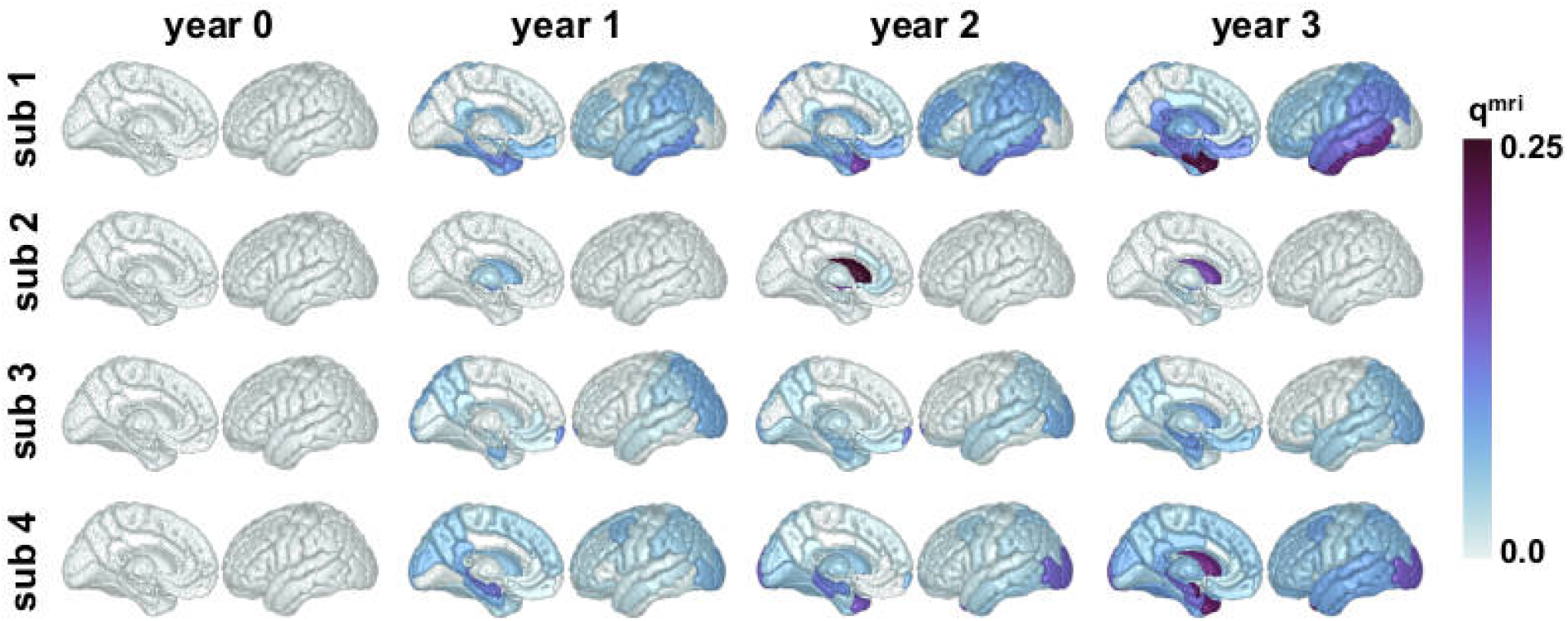
Subject data. Regional atrophy *q*^mri^ normalized to baseline scan illustrated on a template brain. Each row summarizes the data for one subject, at baseline year 0 and three follow up scans.

### 5.2 Posterior distributions

Figure 5 summarizes the personalized posterior distributions for the parameters of our coupled tau-atrophy model. All distributions have clearly moved from the weakly informative prior distributions, indicating successful inference. The estimates for measurement noise in the atrophy data *σ*_a_ are slightly higher than the estimated noise in the tau data *σ*_t_, especially for subjects 1 and 2. The inferred transport coefficients *ρ* for subjects 1,2, and 4 are close to zero. Subject 3 seems to exhibit more pronounced tau dynamics, with an elevated transport coefficient *ρ* and growth coefficient *α* compared to the rest of the cohort. Subject 4 represents an outlier compared to the other subjects when examining the inferred growth rate *α*. The tau data from Subject 4 indicates a negative growth coefficient, which implies that there is more clearance than production of misfolded tau. The inferred tau-induced atrophy coefficients *G*_c_ are of similar magnitude for all subjects.

**Figure 5:**
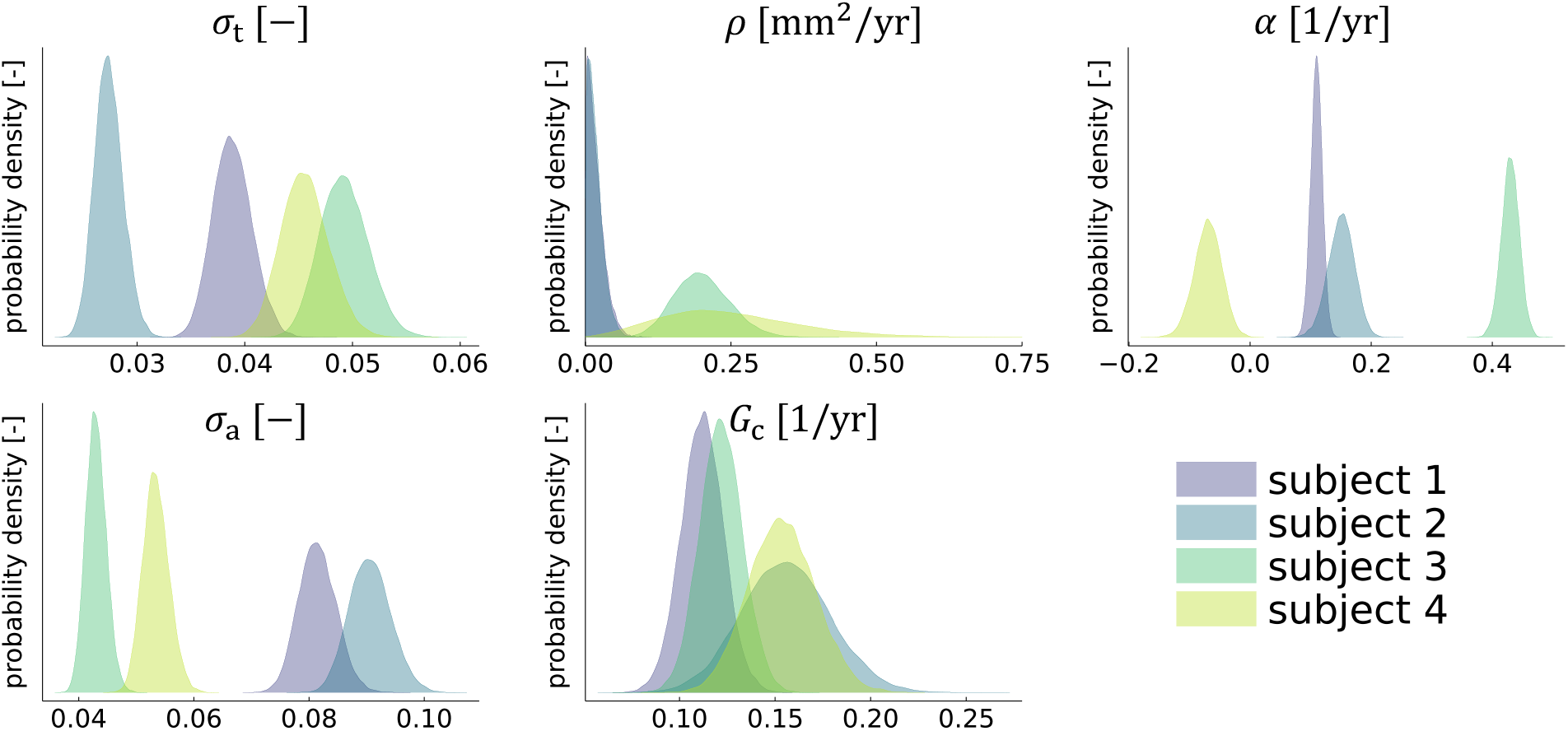
Posterior distributions. Personalized posterior distributions for measurement noise in tau data *σ*_t_, and volume data *σ*_a_, the transport coefficient *ρ*, the growth coefficient *α*, and the tau-induced atrophy coefficient *G_c_*.

### 5.3 Posterior predictive simulations

Figure 6 shows the long- and short-term evolutions of global tau and atrophy predicted by our model for the four examined subjects. The tau and atrophy dynamics are best captured by our model for subject 3. We note characteristic sigmoidal curves for normalized global tau and atrophy in the long-term predictions. All four data points for tau and atrophy lie either within or very close to the 95% credible intervals. Our model captures the dynamics of subject 1 similarly well. Only one of the tau data points lies outside the 95% credible interval, all atrophy data are consistent with the model predictions. When looking at the long-term predictions, the subject exhibits almost linear tau and atrophy dynamics, an observation that is consistent with fairly low transport and growth coefficients inferred for this individual, as discussed in Section 5.2. Our model does a relatively poor job in capturing the dynamics of subjects 2 and 4. For subject 2, three out of four tau data points and two out of four atrophy data points lie outside the 95% credible intervals. By design, our model cannot account for non-monotonic increases or decreases in tau data; it always predicts either an overall increasing or an overall decreasing trend for tau. Our model assumptions imply an increase in atrophy whenever the local tau concentration is positive, which results in an overall increase in longitudinal atrophy for subject 2, even though the observed atrophy does not seem to increase much over time. For subject 4, our model infers an overall declining trend in the global tau concentration from the tau data, again unable to capture the non-monotonically increasing and decreasing data. Consequently, three out of four tau data points lie outside the 95% credible interval. Even though the observed trend in global atrophy data is overall increasing, the inferred negative slope in the tau curve results in a decelerating predicted atrophy rate over time, with whole brain atrophy projected to level out around a value of 25% after 30 years.

**Figure 6:**
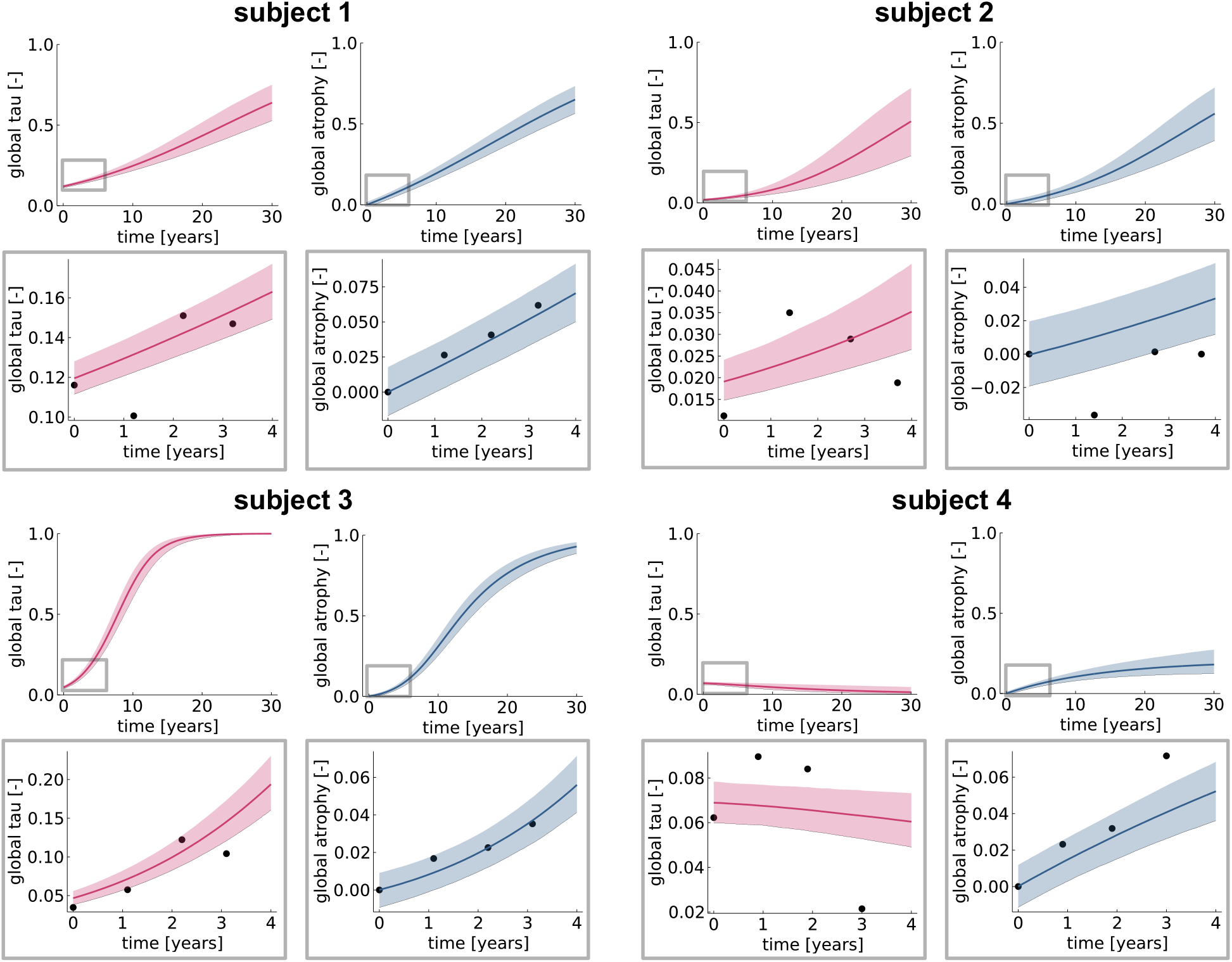
Posterior predictive modeling. Long- and short-term model predictions for four subjects. Each subplot shows the globally averaged tau concentration in red or globally averaged atrophy in blue over time. Gray windows contain detailed views of the first four years of simulation with data points indicated by black circles. Shaded areas around the curves indicate 95% credible intervals.

### 5.4 Personalized atrophy rates

We used our posterior predictive simulations to extract predicted time-dependent atrophy rates for the whole brain, the hippocampus, and the entorhinal cortex in each subject. Initial and maximal atrophy rates are presented in Table 3 and may be compared to the observed rates in the literature summarized in Table 1. Figure 7 shows atrophy rates plotted as a function of time. Subjects 1,2, and 3 exhibit atrophy rates that first increase over time, reach a maximum value, and then decline. We see consistently higher atrophy rates in the hippocampus and the entorhinal cortex than on average across the whole brain, an observation that is in line with the clinical studies summarized in Table 1. The predicted atrophy rates for subject 1 and subject 2 are in a similar range, while those predicted rates for subject 3 are almost twice as high. Subject 4 exhibits different atrophy dynamics than the rest of the cohort, due to the negative inferred growth coefficient, indicating overall declining tau concentration. The initial atrophy rates inferred for subject 4 are in a similar range as those for subject 1 and subject 2.

**Figure 7:**
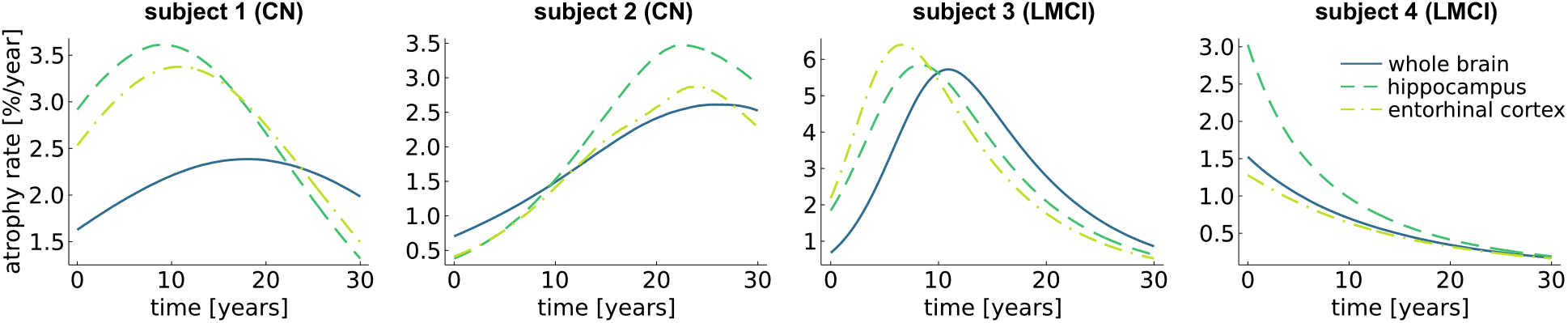
Personalized atrophy rates. Median time-dependent atrophy rates for the whole brain, the hippocampus, and the entorhinal cortex as predicted by our model for each subject.

**Table 3:** Personalized atrophy rates. Atrophy rates inferred by our model at time of baseline scan *t*_0_ and maximum atrophy rates inferred for each subject across whole brain, in the hippocampus, and in the entorhinal cortex.

|  |  | predicted atrophy rates: median (95% CI) in %/year |  |  |  |
| --- | --- | --- | --- | --- | --- |
|  |  | subject 1 | subject 2 | subject 3 | subject 4 |
| whole brain | $t_0$ | 1.63 (1.3-2.0) | 0.71 (0.5-0.9) | 0.68 (0.5-0.9) | 1.53 (1.2-1.9) |
|  | max | 2.39 (2.0-2.8) | 2.61 (1.8-3.4) | 5.72 (5.1-6.3) | 1.53 (1.2-1.9) |
| hippocampus | $t_0$ | 2.93 (1.2-3.9) | 0.39 (0.0-1.0) | 1.84 (1.0-2.8) | 3.03 (2.0-4.4) |
|  | max | 3.62 (3.0-4.4) | 3.47 (1.8-4.6) | 5.85 (5.2-6.6) | 3.03 (2.0-4.4) |
| entorhinal cortex | $t_0$ | 2.53 (1.8-3.5) | 0.41 (0.0-1.0) | 2.18 (1.3-3.1) | 1.26 (0.4-2.4) |
|  | max | 3.37 (2.6-4.1) | 2.94 (0.7-4.5) | 6.4 (5.5-7.2) | 1.26 (0.4-2.4) |

## 6 Discussion

We summarized the characteristics and dynamics of tissue atrophy in Alzheimer’s disease and, from the observed trends, proposed a mathematical model that couples misfolded tau propagation and growth with a model of brain atrophy. We performed a Bayesian analysis to personalize our model parameters from longitudinal multi-model imaging data. Posterior predictions allowed us to examine how tau pathology and brain atrophy may develop over three decades in four examined subjects from the Alzheimer’s Disease Neuroimaging Initiative. From the posterior predictions, we computed time-dependent annual atrophy rates for each subject and compared these personalized atrophy rates with the values reported in the literature.

The inferred tau-induced atrophy coefficient *G_c_*, quantifying the effect of local tau on local tissue atrophy, was similar across subjects despite the wide spectrum of observed tau and atrophy dynamics. On average, we found a median tau-induced atrophy coefficient of *G_c_* = 0.13/year (95%CI: 0.097 – 0.189). The consistency in this parameter is a promising sign towards the ability of accurately predicting personalized future atrophy from tau PET images. Developing a predictive model of tau and atrophy dynamics may allow for advances in personalized estimations of disease progression from a set of tau PET images, or estimations of when certain brain functions might be impaired in an individual patient.

The model given by Equations 4 and 5 reflects key elements of atrophy dynamics discussed in Section 2, and predicts realistic atrophy rates that are consistent with the values reported in the literature for the whole brain, the hippocampus, and the entorhinal cortex. The whole brain atrophy rates in the literature, as summarized in Table 1 range from 0 (0.57) %/year to 0.8 (0.5) %/year for control or cognitively normal subjects and from 0.6 (0.7) %/year to 2.8 (0.5) %/year for subjects with mild cognitive impairment or Alzheimer’s disease. Reported hippocampal atrophy rates lie between 0.86 (1.27) %/year and 3.3 (2.4) %/year for cognitively normal subjects and between 2.39 (2.26) %/year and 5.19 (0.75) %/year for cognitively impaired patients. For the entorhinal cortex, the reported values range from 0.52 (1.13) %/year to 5.1 (5.1) %/year for cognitively normal and from 1.93 (1.56) %/year to 8.4 (9.2) %/year for cognitively impaired subjects. Apart from subject 4, the plots in Figure 7 show atrophy rates first increasing, and then decreasing over time, consistent with the observations of early atrophy acceleration and late atrophy deceleration. Additionally, we see consistently higher atrophy rates in the hippocampus and the entorhinal cortex than on average across the whole brain. These two regions are known to be affected by tau and atrophy early in the disease, and their atrophy rates are often used as biomarkers for early diagnosis. Thus, it makes sense that atrophy rates in the hippocampus and entorhinal cortex are higher than average and tend to reach their peak rate earlier than the whole brain on average. For subject 1, the predicted maximum atrophy rates are in good alignment with the rates observed in the literature for patients with Alzheimer’s disease or mild cognitive impairment. For a cognitively normal subject, the current diagnosis for subject 1, the starting atrophy rates are on the higher end compared to the literature, indicating a potential conversion to mild cognitive impairment in the future. This is plausible due to the positive amyloid status associated with subject 1. Subject 2 exhibits slightly lower beginning atrophy rates, but similar maximum rates as subject 1, which are also comparable with the literature. For subject 3, our model predicts a steep increase in atrophy rates over the first 10 years after the baseline scan. The predicted maximum whole brain atrophy rate is higher than what is observed on average in the literature, even for Alzheimer’s with fast cognitive decline and early onset. The maximum hippocampal and entorhinal atrophy rates are on the high end, but within the range of observed values. Our model predicts declining atrophy rates for subject 4, starting at about 1.5 %/year for whole brain and entorhinal atrophy rate, and about 3 %/year for the hippocampal atrophy rate, and subsequently declining to zero after around 30 years. This trend does not reflect the observations of accelerating atrophy rates described in Section 2, unless this subject has already progressed so far in the disease, that we are only observing the decline in atrophy rates that is described for very late disease stages. The beginning atrophy rates are well in the range of values reported in the literature for whole brain, hippocampus and entorhinal cortex.

This study comes with a number of limitations. First, we constrained our model calibration to a very small, preliminary cohort of four subjects. However, our work demonstrates the feasibility of inferring model parameters from multi-modal data and will be expanded to more subjects to test for the overall statistical significance of our preliminary results. Second, the four examined subjects produced a wide range of tau and atrophy dynamics. Our model does well in explaining the data of subjects 1 and 3 while the results for subjects 2 and 4 suggest that accounting for possible non-monotony in Equation 4 may further improve the descriptive capacity of the model. Nevertheless, observing the spectrum of possible dynamics and applicability of the model to even a small preliminary data sample is valuable and highlights the challenges that might be encountered in a larger study. A third limitation is posed by longitudinal data availability. In particular, our data was limited to four time points due to the relative novelty and limited availability of tau PET data in the ADNI database. Specifically, out of the 2492 study participants with imaging data in ADNI, each has on average 4.57 ± 3.318 structural MRI scans, but only 0.475 ± 0.826 tau PET scans. Only 786 participants have tau PET data at all, and out of those each subject has on average 1.506 ± 0.782 longitudinal scans, with a minimum of one scan and maximum of five scans per subject. As tau PET becomes a more established imaging modality across participating study centers, the amount of available longitudinal data will naturally increase, allowing us to improve our model evaluations and assess its predictive capabilities. Longitudinal data availability also constrains the number of model parameters that can be reliably inferred while avoiding overfitting. This is an argument for selecting model formulations that balance model expressiveness with data concerns. For instance, the atrophy model presented in Equation 5 does not account for age-related atrophy in the absence of misfolded tau. However, as mentioned in Section 2, it is well established that some amount of tissue atrophy occurs during normal ageing without the presence of significant tau pathology. Limited data availability also prompted us to exclude any attempt to infer deviations in initial conditions, extracted from the baseline tau PET scan, as a variable in our inference approach. Such an approach would introduce new distributions to be inferred, at every cortical region. With future increases in longitudinal data, our model and inference approach could be augmented to reflect more complex atrophy characteristics and take into account potential deviations in initial conditions.

Bayesian methods inherently facilitate working with continuously updated data, a feature that will make it easy to seamlessly update the personalizations with more time points becoming available in the future. In upcoming work, we plan to investigate the identifiability of more complex tau-atrophy models, taking into account the limited amount and inherent noisiness of available imaging data for calibration.

## 7 Conclusion

We presented a high-level overview of what we know, today, about brain atrophy in Alzheimer’s disease and demonstrated how we can translate this knowledge into a computational model. Our model captures a coupling between tau pathology and atrophy, and naturally features an early acceleration and late deceleration of atrophy rates. This is made possible through a direct dependence on the local concentration of pathological tau and a saturation term. Personalizing our model with a small data set using a probabilistic approach gives us an initial idea of the ranges of model parameters and associated uncertainties that might occur in real subject data. Strikingly, the magnitude of the tau-induced atrophy coefficient, a direct indicator for the correlation of misfolded tau and tissue atrophy, is fairly consistent across all subjects with a median value of 0.13/year. Using our personalized model, we created long-term predictions for the tau and atrophy dynamics in four subjects and discussed the performance of our model for each individual scenario. We observed that our model generates realistic predictions of atrophy rates that agree well with the rates reported in the literature. Once comprehensively calibrated with a larger set of longitudinal data, our model can be used as a personalized diagnostic and predictive tool to estimate future atrophy from tau PET recordings or serve as a comparative control in treatment studies.

## Acknowledgements

The work of E. Kuhl was supported by the National Science Foundation grant CMMI 1727268. The work of A. Goriely was supported by the Engineering and Physical Sciences Research Council grant EP/R020205/1. The work of A. Schaefer was supported by a Brit and Alex d’Arbeloff Stanford Graduate Fellowship to AS. The work of P. Chaggar was supported by funding from the Engineering and Physical Sciences Research Council grant EP/L016044/1 and Roche. The work of T. Thompson was supported partially the John Fell Oxford University Press Research Fund grant 000872 (project code BKD00160) to TT, and partially by the Engineering and Physical Sciences Research Council grant EP/R020205/1 to AG.

Data collection and sharing for this project was funded by the Alzheimer’s Disease Neuroimaging Initiative (ADNI) (National Institutes of Health Grant U01 AG024904) and DOD ADNI (Department of Defense award number W81XWH-12-2-0012). ADNI is funded by the National Institute on Aging, the National Institute of Biomedical Imaging and Bioengineering, and through generous contributions from the following: AbbVie, Alzheimer’s Association; Alzheimer’s Drug Discovery Foundation; Araclon Biotech; BioClinica, Inc.; Biogen; Bristol-Myers Squibb Company; CereSpir, Inc.; Cogstate; Eisai Inc.; Elan Pharmaceuticals, Inc.; Eli Lilly and Company; EuroImmun; F. Hoffmann-La Roche Ltd and its affiliated company Genentech, Inc.; Fujirebio; GE Healthcare; IXICO Ltd.; Janssen Alzheimer Immunotherapy Research & Development, LLC.; Johnson & Johnson Pharmaceutical Research & Development LLC.; Lumosity; Lundbeck; Merck & Co., Inc.; Meso Scale Diagnostics, LLC.; NeuroRx Research; Neurotrack Technologies; Novartis Pharmaceuticals Corporation; Pfizer Inc.; Piramal Imaging; Servier; Takeda Pharmaceutical Company; and Transition Therapeutics. The Canadian Institutes of Health Research is providing funds to support ADNI clinical sites in Canada. Private sector contributions are facilitated by the Foundation for the National Institutes of Health (www.fnih.org). The grantee organization is the Northern California Institute for Research and Education, and the study is coordinated by the Alzheimer’s Therapeutic Research Institute at the University of Southern California. ADNI data are disseminated by the Laboratory for Neuro Imaging at the University of Southern California.

